# Genome sequencing and analysis of two early-flowering cherry (*Cerasus* × *kanzakura*) varieties, ‘Kawazu-zakura’ and ‘Atami-zakura’

**DOI:** 10.1101/2021.09.08.459382

**Authors:** Kenta Shirasawa, Akihiro Itai, Sachiko Isobe

## Abstract

To gain genetic insights into the early-flowering phenotype of ornamental cherry, also known as sakura, we determined the genome sequences of two early-flowering cherry (*Cerasus* × *kanzakura*) varieties, ‘Kawazu-zakura’ and ‘Atami-zakura’. Since the two varieties are interspecific hybrids, likely derived from crosses between *Cerasus campanulata* (early-flowering species) and *Cerasus speciosa*, we employed the haplotype-resolved sequence assembly strategy. Genome sequence reads obtained from each variety by single molecule real-time sequencing (SMRT) were split into two subsets, based on the genome sequence information of the two probable ancestors, and assembled to obtain haplotype-phased genome sequences. The resultant genome assembly of ‘Kawazu-zakura’ spanned 519.8 Mb with 1,544 contigs and an N50 value of 1,220.5 kb, while that of ‘Atami-zakura’ totaled 509.6 Mb with 2,180 contigs and an N50 value of 709.1 kb. A total of 72,702 and 72,528 potential protein-coding genes were predicted in the genome assemblies of ‘Kawazu-zakura’ and ‘Atami-zakura’, respectively. Gene clustering analysis identified 2,634 clusters uniquely presented in the *C. campanulata* haplotype sequences, which might contribute to its early-flowering phenotype. Genome sequences determined in this study provide fundamental information for elucidating the molecular and genetic mechanisms underlying the early-flowering phenotype of ornamental cherry tree varieties and their relatives.

## Introduction

Flowering cherry, called sakura in Japanese, is an ornamental plant popular worldwide. A major *Cerasus* × *yedoensis* cultivar ‘Somei-Yoshino’, which is an interspecific hybrid of *Cerasus spachiana* and *Cerasus speciosa*^1^, usually blooms from March to April in Japan. In addition, early-flowering sakura species, such as *Cerasus campanulata*, usually bloom 1–2 months earlier than ‘Somei-Yoshino’, and its interspecific hybrids such as *Cerasus* × *kanzakura* also exhibit early flowering. *C*. × *kanzakura* is considered a hybrid between *C. campanulata* and *Cerasus speciosa* and/or *Cerasus jamasakura*^2^, but its origin is still debated. Two *C*. × *kanzakura* cultivars, ‘Kawazu-zakura’ and ‘Atami-zakura’, also bloom early (January and February, respectively); however, the molecular mechanisms underlying their early-flowering phenotype remain unknown. Although the mechanisms of early flowering in Rosaceae family members, Japanese plum (*Prunus mume*) and peach (*Prunus persica*), which flower in February and March, respectively, are well known^3^, it remains unclear whether these mechanisms are common between *Cerasus* and *Prunus*.

Genome sequence analysis provides information on nucleotide polymorphisms and gene copy number variation, which can lead to phenotypic differences among individuals and cultivars^4^. Pan-genomics, which involves de novo genome sequencing of multiple lines within a species, is conducted to obtain information on variation in all genes within a species to understand the origin of the organism under study^5,6^. In ‘Somei-Yoshino’, haplotype-phased genome sequences have been reported, and comprehensive changes in gene expression during floral bud development that contribute toward flowering have been revealed by time-course transcriptome analysis^7^. Therefore, comparative genomics of multiple lines of flowering cherry varieties, such as ‘Kawazu-zakura’, ‘Atami-zakura’, and ‘Somei-Yoshino’, could provide genetic insights into their early-flowering phenotypes.

A trio-binning strategy^8^, previously used in a bovine F1 hybrid to resolve two haplotype-phased genome sequences, was recently applied to ‘Somei-Yoshino’^7^. Genes associated with the early-flowering phenotype of ‘Kawazu-zakura’ and ‘Atami-zakura’ were assumed to be encoded by the *C. campanulata* haplotype sequences. Therefore, in this study, we used the trio-binning strategy to determine the haplotype-phased sequences of ‘Kawazu-zakura’ and ‘Atami-zakura’. Comparative analysis of three sakura genomes (‘Kawazu-zakura’, ‘Atami-zakura’, and ‘Somei-Yoshino’) facilitated the identification of genes unique to the *C. campanulata* haplotype sequences of ‘Kawazu-zakura’ and ‘Atami-zakura’ as candidates responsible for the early-flowering phenotype of these varieties.

## Materials and methods

### Plant materials and DNA extraction

Two early-flowering cherry (*Cerasus* × *kanzakura*) varieties, ‘Kawazu-zakura’ and ‘Atami-zakura’, were used in this study. Both varieties were planted at the orchard of Kyoto Prefectural University (Kyoto, Japan). Genome DNA was extracted from young leaves by a modified sodium dodecyl sulfate (SDS) method^9^.

### Genome size estimation

Software tools used for data analyses are listed in Supplementary Table S1. Genome libraries for short-read sequencing were prepared with the TruSeq DNA PCR-Free Sample Prep Kit (Illumina, San Diego, CA, USA), and sequenced on the NextSeq 500 platform (Illumina, San Diego, CA, USA) in paired-end, 150 bp mode. The genome size was estimated with Jellyfish.

### De novo genome sequence assembly and reference-guided contig ordering and orientation

Genomes of the two cherry varieties were sequenced using the single molecule real-time (SMRT) sequencing technology. Long-read DNA libraries were constructed using the SMRTbell Express Template Prep Kit 2.0 (PacBio, Menlo Park, CA, USA) and sequenced on SMRT cells (1M v3 LR) in a PacBio Sequel system (PacBio). Raw sequence reads of each variety were divided into two subsets with the trio-binning strategy^8^ using the short-read data of *C. campanulata* (‘Kanhi-zakura’) and *C. speciosa* (‘Ohshima-zakura’) (DDBJ sequence archive accession no.: DRA008096)^7^. The sequence read subsets were assembled separately with Falcon or Canu to build haplotype-phased diploid genome sequences. Sequence errors in the contigs were corrected twice using long reads with ARROW. Potential contaminating sequence reads from organelle genomes were identified by alignments with the chloroplast and mitochondrial genome sequences of *Prunus avium* (GenBank accession no.: MK622380 and MK816392) with Minimap2, and then removed from the final assemblies. Haplotype-phased sequences, based on binning with *C. campanulata* and *C. speciosa*, were aligned against the *C. spachiana* and *C. speciosa* haplotype sequences, respectively, of the ‘Somei-Yoshino’ genome using Ragoo to build pseudomolecule sequences. Genome sequences were compared with D-Genies.

### Gene prediction and repetitive sequence analysis

Potential protein-coding genes were predicted with the MAKER pipeline, which was based on peptide sequences predicted from the genome sequences of sweet cherry (PAV_r1.0)^10^, peach (v2.0.a1)^11^, and Japanese plum^12^. Short genes (<300 bp) as well as genes predicted with an annotation edit distance (AED) greater than 0.5, which is proposed as a threshold for good annotations in the MAKER protocol, were removed to facilitate the selection of high-confidence (HC) genes. Functional annotation of the predicted genes was performed with Hayai-Annotation Plants. Gene clustering was performed with OrthoFinder.

Repetitive sequences in the pseudomolecules were identified with RepeatMasker using repeat sequences registered in Repbase and a *de novo* repeat library built with RepeatModeler. The identified repetitive sequences were classified into nine types, in accordance with RepeatMasker: short interspersed nuclear elements (SINEs), long interspersed nuclear elements (LINEs), long terminal repeat (LTR) elements, DNA elements, small RNAs, satellites, simple repeats, low complexity repeats, and unclassified.

## Results and data description

### De novo assembly of ‘Kawazu-zakura’ and ‘Atami-zakura’ genomes

Short reads amounting to 64.0 and 127.7 Gb were obtained for ‘Kawazu-zakura’ and ‘Atami-zakura’, respectively. The genome sizes of ‘Kawazu-zakura’ and ‘Atami-zakura’ were estimated at 672.7 and 675.2 Mb, respectively (Figure 1). Since ‘Kawazu-zakura’ and ‘Atami-zakura’ are interspecific hybrids, we used a trio-binning strategy to establish haplotype-resolved genome assemblies representing each parental genome sequence.

**Figure 1.**
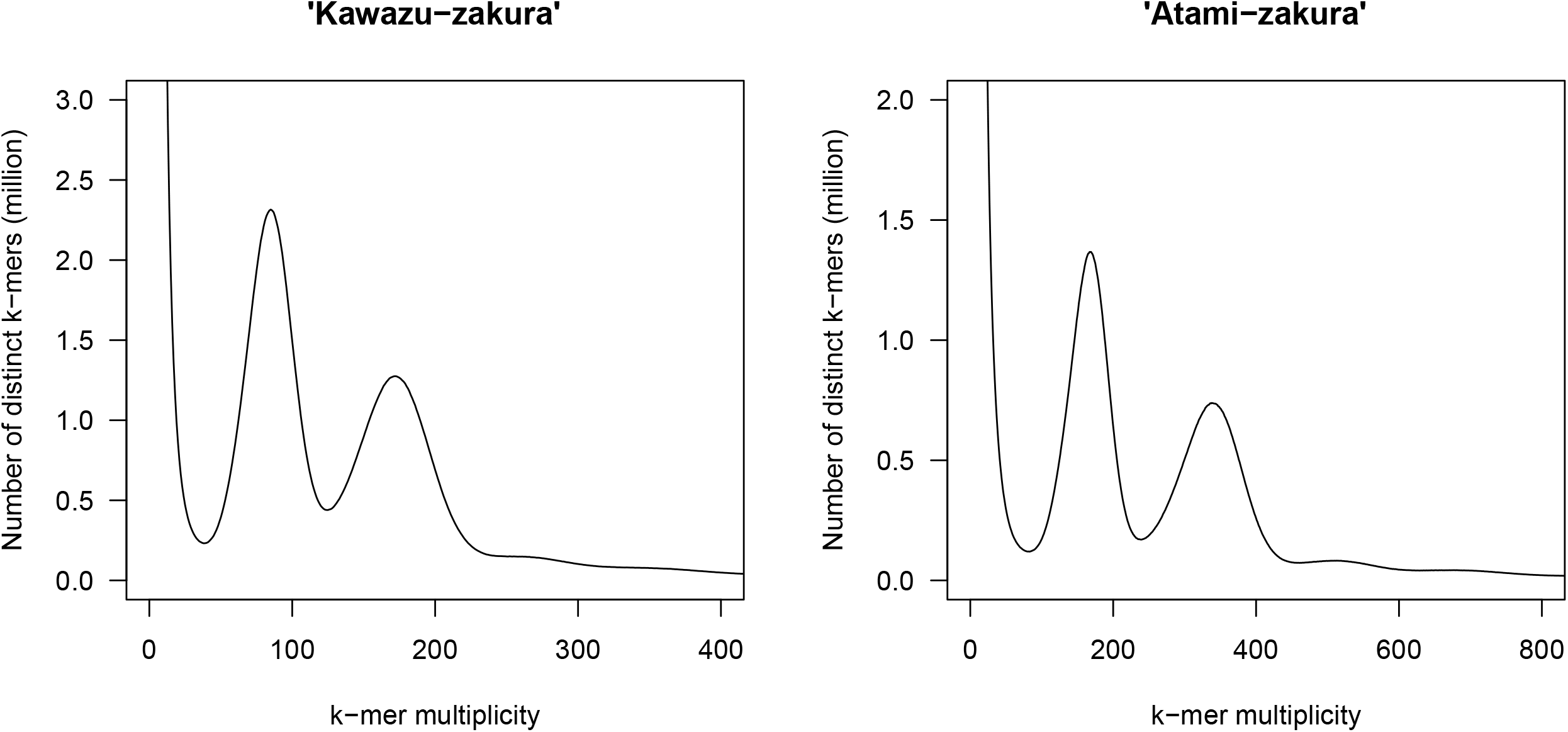
Estimation of the genome size of two flowering cherry (*Cerasus* × *kanzakura*) varieties, ‘Kawazu-zakura’ and ‘Atami-zakura’, based on *k*-mer analysis (*k* = □17), with the given multiplicity values.

Long-read data (34.9 Gb) of ‘Kawazu-zakura’ obtained from two SMRT cells were divided into two subsets (17.2 and 17.6 Gb), in accordance with the short-read data of potential parental species, *C. campanulata* and *C. speciosa*^7^, respectively. Reads in each subset were independently assembled with Falcon to construct contigs representing the two haplotype sequences. Potential errors in the haplotype sequences were corrected with long reads, and sequences of organelle genomes were removed to obtain the final assembly of the diploid genome of ‘Kawazu-zakura’. The resulting assemblies consisted of *C. campanulata* (262.2 Mb, N50 = 1.4 Mb) and *C. speciosa* (257.6 Mb, N50 = 1.1 Mb) haplotypes (Table 1), and were designated as KWZcam_r1.0 and KWZspe_r1.0, respectively. Although the total assembly size was shorter than the estimated size, the complete BUSCO scores of KWZcam_r1.0 and KWZspe_r1.0 were 93.1% and 96.7%, respectively, indicating that the assemblies were complete (Table 1). The two assemblies were merged to generate KWZ_r1.0, with a complete BUSCO score of 98.0%.

**Table 1.** Statistics of the contig sequences of two flowering cherry (*Cerasus* × *kanzakura*) cultivars, ‘Kawazu-zakura’ and ‘Atami-zakura’

|  | KWZ_r1.0 | KWZcam_r1.0 | KWZspe_r1.0 | ATM_r1.0 | ATMcam_r1.0 | ATMspe_r1.0 |
| --- | --- | --- | --- | --- | --- | --- |
| Total contig size (bases) | **519,843,677** | 262,196,010 | 257,647,667 | **509,633,549** | 267,393,285 | 242,240,264 |
| Number of contigs | **1,544** | 783 | 761 | **2,180** | 1,124 | 1,056 |
| Contig N50 length (bases) | **1,220,495** | 1,445,144 | 1,108,133 | **709,113** | 853,547 | 569,444 |
| Longest contig size (bases) | **8,019,066** | 5,955,677 | 8,019,066 | **5,799,312** | 5,799,312 | 3,381,444 |
| Gap (bases) | **0** | 0 | 0 | **0** | 0 | 0 |
| Complete BUSCOs | **98.2%** | 93.1% | 96.7% | **98.0%** | 93.4% | 93.5% |
| Single-copy BUSCOs | **7.5%** | 86.7% | 89.0% | **16.0%** | 86.5% | 87.8% |
| Duplicated BUSCOs | **90.7%** | 6.4% | 7.7% | **82.0%** | 6.9% | 5.7% |
| Fragmented BUSCOs | **0.3%** | 0.7% | 0.4% | **0.4%** | 0.7% | 1.6% |
| Missing BUSCOs | **1.5%** | 6.2% | 2.9% | **1.6%** | 5.9% | 4.9% |
| #Genes | **72,702** | 36,281 | 36,421 | **72,528** | 36,264 | 36,264 |

The ‘Atami-zakura’ genome was sequenced in parallel with the ‘Kawazu-zakura’ genome. Long-read data of ‘Atami-zakura’ (14.3 Gb) were obtained from two SMRT cells, and divided into two subsets (7.4 and 6.8 Gb) using the short-read data of *C. campanulata* and *C. speciosa*^7^, respectively. The reads were assembled with Falcon to generate two haplotype contig sequences. However, the sizes of the two assemblies (139.9 and 110.0 Mb) were much smaller than the estimated sizes. Therefore, we used Canu to obtain long assemblies. This was followed by potential sequence error correction and organelle genome sequence removal. The sizes of the resultant assemblies were improved to 267.4 Mb (N50 = 853.5 kb) and 242.2 Mb (N50 = 569.4 Mb) for the *C. campanulata* and *C. speciosa* haplotypes, respectively (Table 1), and the assemblies were designated as ATMcam_r1.0 and ATMspe_r1.0, respectively. The complete BUSCO scores were 93.4% and 93.5% for ATMcam_r1.0 and ATMspe_r1.0, respectively (Table 1), and 98.2% for the merged assembly (ATM_r1.0).

### Reference-guided pseudomolecule sequence construction

Since the genome structures are well conserved across the *Cerasus* and *Prunus* species^7^, we used the two haplotype pseudomolecule sequences of the ‘Somei-Yoshino’ genome, CYEspachiana_r3.1 and CYEspeciosa_r3.1, as references to establish the pseudomolecule sequences of ‘Kawazu-zakura’ and ‘Atami-zakura’. A total of 777 and 746 contigs of KWZcam_r1.0 and KWZspe_r1.0, respectively, were aligned against CYEspachiana_r3.1 and CYEspeciosa_r3.1 sequences, respectively. The lengths of the resultant ‘Kawazu-zakura’ pseudomolecule sequences were 256.7 Mb (KWZcam_r1.0) and 246.5 Mb (KWZspe_r1.0) (Table 2). On the other hand, 1,110 ATMcam_r1.0 and 1,041 ATMspe_r1.0 contigs were aligned with the CYEspachiana_r3.1 and CYEspeciosa_r3.1 sequences, respectively, and the lengths of the ‘Atami-zakura’ pseudomolecule sequences obtained were 261.5 Mb (ATMcam_r1.0) and 238.9 Mb (ATMspe_r1.0) (Table 2). The pseudomolecule sequences of ‘Kawazu-zakura’ and ‘Atami-zakura’ genomes covered the entire genome sequence of ‘Somei-Yoshino’ (Figure 2).

**Table 2.**
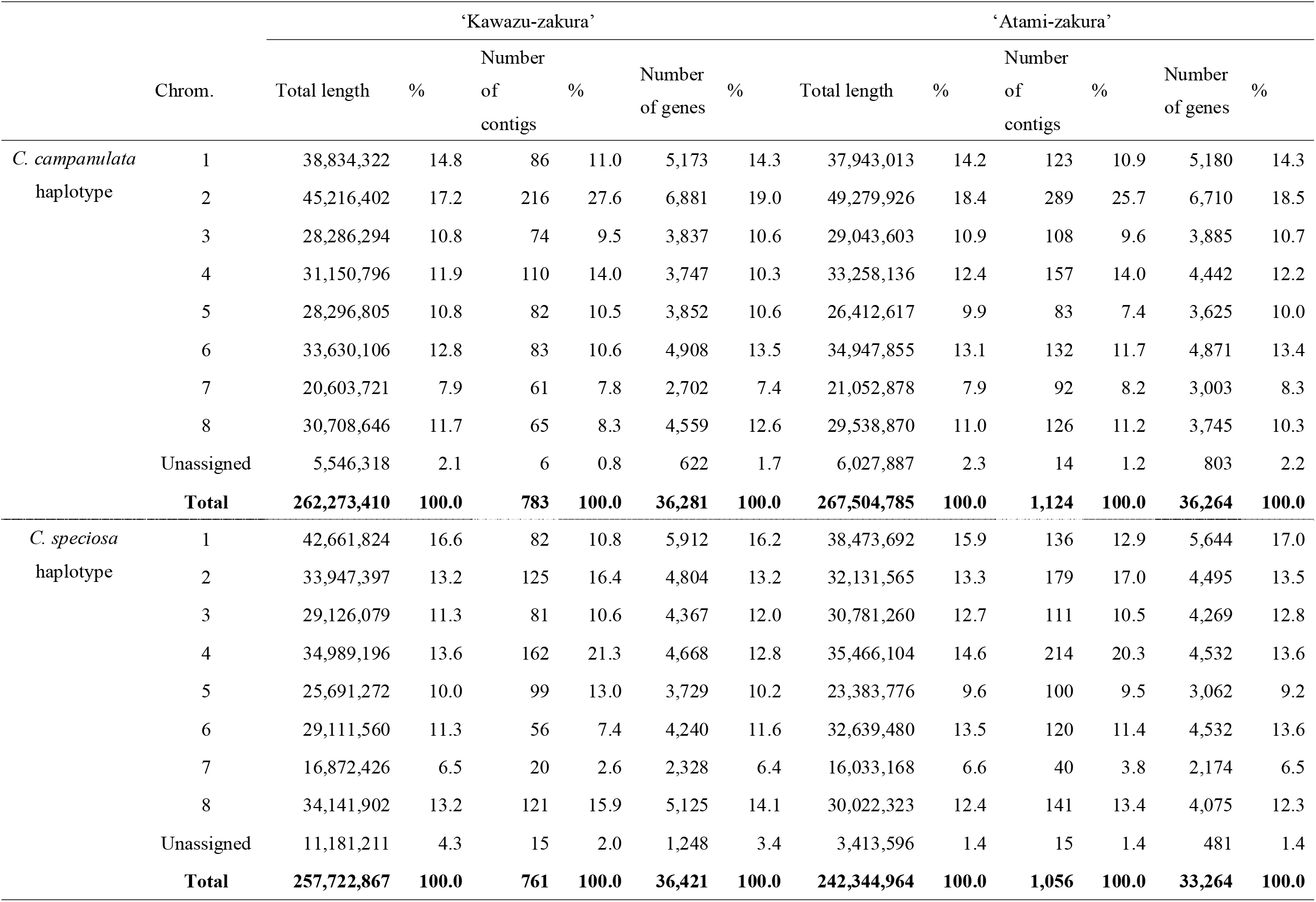
Statistics of the pseudomolecule sequences of flowering cherry (*C*. × *kanzakura*) cultivars, ‘Kawazu-zakura’ and ‘Atami-zakura’

**Figure 2.**
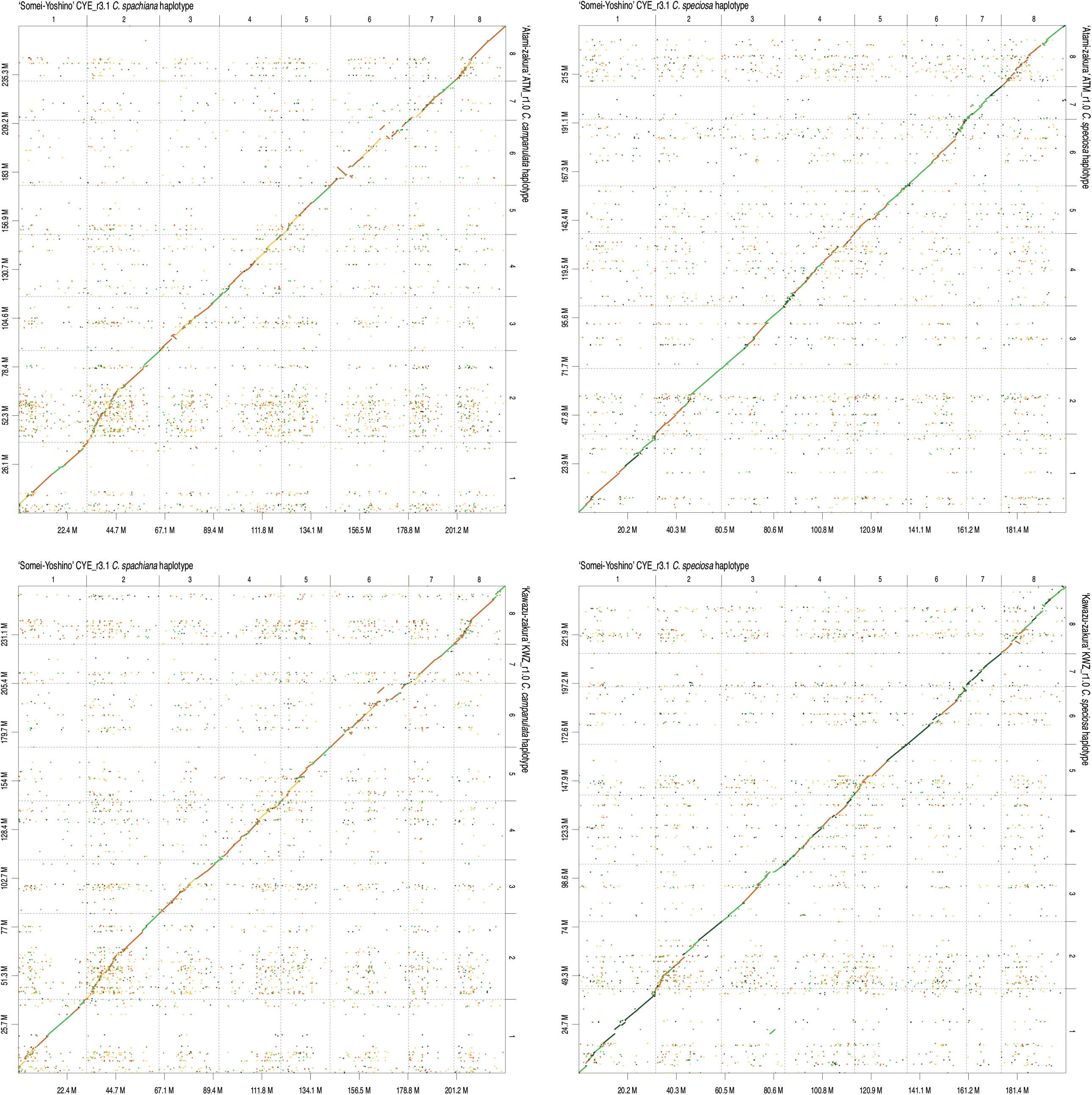
Comparative analysis of the genome sequence and structure of flowering cherry varieties, ‘Atami-zakura’, ‘Kawazu-zakura’, and ‘Somei-Yoshino’. Chromosome numbers are indicated above the x-axis and on the right side of the y-axis. Genome sizes (Mb) are below the x-axis and on the left side of the y-axis.

### Gene and repetitive sequence predictions

A total of 36,264 and 36,264 HC protein-coding genes were predicted in KWZcam_r1.0 and KWZspe_r1.0 assemblies, respectively (Table 2). The complete BUSCO scores of genes in the KWZcam_r1.0 and KWZspe_r1.0 were 88.3% and 86.6%, respectively, while the BUSCO score of all 72,582 genes was 97.0%. Functional gene annotation revealed that 9,430, 17,907, and 12,603 sequences were assigned to Gene Ontology (GO) slim terms in the biological process, cellular component, and molecular function categories, respectively, and 2,264 genes had enzyme commission numbers.

On the other hand, 36,281 and 36,421 HC genes were predicted in ATMcam_r1.0 and ATMspe_r1.0 assemblies, respectively (Table 2). Complete BUSCOs of genes in ATMcam_r1.0 and ATMspe_r1.0 were 88.3% and 86.6%, respectively, while that of all 72,702 genes was 96.8%. According to the functional gene annotation, 9,836, 18,586, and 13,020 sequences were assigned to GO slim terms in the biological process, cellular component, and molecular function categories, respectively, and 2,301 genes had enzyme commission numbers.

Repeat sequences occupied varying proportions of the different genome assemblies: 48.0% (KWZcam_r1.0), 45.7% (KWZspe_r1.0), 47.7% (ATMcam_r1.0), and 43.2% (ATMspe_r1.0). LTR elements were the most abundant repetitive sequences (15.1–17.7%), followed by unclassified repeats (12.7–13.7%) and DNA transposons (11.1–13.2%) (Table 3).

**Table 3.**
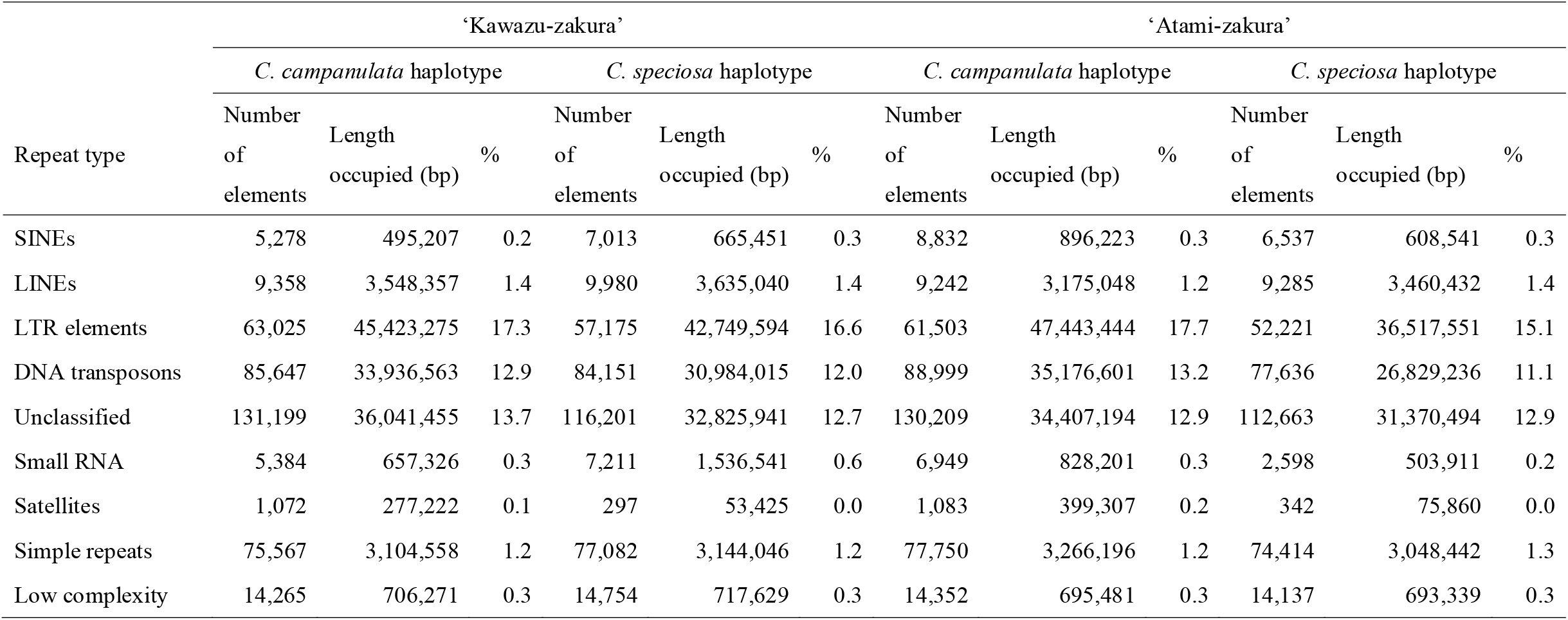
Repetitive sequences in two flowering cherry (*C*. × *kanzakura*) cultivars, ‘Kawazu-zakura’ and ‘Atami-zakura’

| Repeat type | ‘Kawazu-zakura’ |  |  |  |  |  | ‘Atami-zakura’ |  |  |  |  |  |
| --- | --- | --- | --- | --- | --- | --- | --- | --- | --- | --- | --- | --- |
|  | *C. campanulata* haplotype |  |  | *C. speciosa* haplotype |  |  | *C. campanulata* haplotype |  |  | *C. speciosa* haplotype |  |  |
|  | Number of elements | Length occupied (bp) | % | Number of elements | Length occupied (bp) | % | Number of elements | Length occupied (bp) | % | Number of elements | Length occupied (bp) | % |
| SINEs | 5,278 | 495,207 | 0.2 | 7,013 | 665,451 | 0.3 | 8,832 | 896,223 | 0.3 | 6,537 | 608,541 | 0.3 |
| LINEs | 9,358 | 3,548,357 | 1.4 | 9,980 | 3,635,040 | 1.4 | 9,242 | 3,175,048 | 1.2 | 9,285 | 3,460,432 | 1.4 |
| LTR elements | 63,025 | 45,423,275 | 17.3 | 57,175 | 42,749,594 | 16.6 | 61,503 | 47,443,444 | 17.7 | 52,221 | 36,517,551 | 15.1 |
| DNA transposons | 85,647 | 33,936,563 | 12.9 | 84,151 | 30,984,015 | 12.0 | 88,999 | 35,176,601 | 13.2 | 77,636 | 26,829,236 | 11.1 |
| Unclassified | 131,199 | 36,041,455 | 13.7 | 116,201 | 32,825,941 | 12.7 | 130,209 | 34,407,194 | 12.9 | 112,663 | 31,370,494 | 12.9 |
| Small RNA | 5,384 | 657,326 | 0.3 | 7,211 | 1,536,541 | 0.6 | 6,949 | 828,201 | 0.3 | 2,598 | 503,911 | 0.2 |
| Satellites | 1,072 | 277,222 | 0.1 | 297 | 53,425 | 0.0 | 1,083 | 399,307 | 0.2 | 342 | 75,860 | 0.0 |
| Simple repeats | 75,567 | 3,104,558 | 1.2 | 77,082 | 3,144,046 | 1.2 | 77,750 | 3,266,196 | 1.2 | 74,414 | 3,048,442 | 1.3 |
| Low complexity | 14,265 | 706,271 | 0.3 | 14,754 | 717,629 | 0.3 | 14,352 | 695,481 | 0.3 | 14,137 | 693,339 | 0.3 |

### Clustering analysis of flowering time related genes in cherry varieties

Four sets of genes predicted in the haplotype-phased genomes of ‘Kawazu-zakura’ and ‘Atami-zakura’ clustered with two sets of genes in the two haploid sequences of ‘Somei-Yoshino’. A total of 35,226 clusters were obtained, of which 10,702 were common across all six gene sets. The early-flowering phenotype of *C*. × *kanzakura* could be explained by genes uniquely present in the *C. campanulata* haplotype sequences. In the *C. campanulata* haplotype sequences of ‘Kawazu-zakura’ and ‘Atami-zakura’ genomes, a total of 2,634 clusters were found to include 3,113 and 3,123 genes, respectively, suggesting that these genes might be associated with the early-flowering phenotype of ‘Kawazu-zakura’ and ‘Atami-zakura’.

## Conclusion and future perspectives

Here, we report haplotype-phased genome assemblies of two early-flowering cherry (*C*. × *kanzakura*) cultivars, ‘Kawazu-zakura’ and ‘Atami-zakura’, both of which are interspecific hybrids derived from *C. campanulata* and *C. speciosa*. Although the origin of *C*. × *kanzakura* remains unclear, *C. campanulata* and *C. speciosa* and/or *C. jamasakura* are considered as its potential parents^2^. Another possibility is that ‘Atami-zakura’ originated from *C. jamasakura* and *C. campanulata*^13^. This is supported by the fact that our attempt to divide the long reads of ‘Atami-zakura’ into two subsets using short-read data of *C. serrulata* (closely related to *C. jamasakura*^7^) and *C. campanulata* failed (data not shown). Therefore, we used short reads of *C. campanulata* and *C. speciosa* for both ‘Kawazu-zakura’ and ‘Atami-zakura’. This result suggests that both ‘Kawazu-zakura’ and ‘Atami-zakura’ are closely related to *C. campanulata* and *C. speciosa*.

Clustering analysis of genes predicted in the genomes of ‘Kawazu-zakura’ and ‘Atami-zakura’ together with those of ‘Somei-Yoshino’ revealed that 2,634 gene clusters were uniquely present in the genome of *C. campanulata* but absent from the genomes of *C. spachiana* and *C. speciosa*. Such copy number variation (or presence/absence variation) of genes could explain the early-flowering phenotype of ‘Kawazu-zakura’ and ‘Atami-zakura’. Previously, we performed a time-course transcriptome analysis of the floral buds and flowers of ‘Somei-Yoshino’ to clarify gene expression patterns during flowering^7^. A similar time-course transcriptome analysis could be applied to ‘Kawazu-zakura’ and ‘Atami-zakura’. Comparative transcriptome analysis of three cultivars could identify the genes responsible for the early-flowering phenotype of sakura. Furthermore, comparative transcriptome analysis of Japanese apricot and peach^3^ could reveal the genetic mechanisms controlling flowering time across all *Prunus* and *Cerasus* species.

Although several flowering cherry cultivars are known to bloom in late-spring, fall, and winter seasons^7^, genome sequences of only a few of these cultivars are publicly available^7,14,15^. Comparative genomics and transcriptomics, also known as pan-genomics^4-6^, of sakura would provide insights into the origins of these cultivars and their flowering mechanisms, which could facilitate the development of new cultivars with attractive flower characteristics and provide us with the ability to forecast the date of sakura blooming.

## Acknowledgments

We thank Y. Kishida, C. Minami, H. Tsuruoka, and A. Watanabe (Kazusa DNA Research Institute) for technical assistance.

## Data availability

Sequence reads are available from the DNA Data Bank of Japan (DDBJ) Sequence Read Archive (DRA) database (accession no.: DRA012553). The DDBJ accession numbers of assembled sequences are BPUM01000001–BPUM01000783 (KWZcam_r1.0), BPUM01000784–BPUM01001544 (KWZspe_r1.0), BPUL01000001–BPUL01001124 (ATMcam_r1.0), and BPUL01001125–BPUL01002180 (ATMspe_r1.0). The genome sequence information generated in this study is available at Plant GARDEN (https://plantgarden.jp).

## Supporting information

**Supplementary Table S1** Software tools used for genome assembly and gene prediction.

